# Cortical area and subcortical volume mediate the effect of parental education and adverse experiences on cognitive performance in youth

**DOI:** 10.1101/160424

**Authors:** Chintan M. Mehta, Jeffrey G. Malins, Kimberly G. Noble, Jeffrey R. Gruen

## Abstract

Early adversity and socioeconomic disadvantage are risk factors associated with diminished cognitive outcomes during development. Recent studies also provide evidence that upbringings characterized by stressful experiences and markers of disadvantage during childhood, such as lower parental education or household income, are associated with variation in brain structure. Although disadvantage often confers adversity, these are distinct risk factors whose differential influences on neurodevelopment and neurocognitive outcomes are not well characterized. We examined pathways linking parental education, adverse experiences, brain structure, and cognitive performances through an analysis of 1,413 typically-developing youth, ages 8 through 21, in the Philadelphia Neurodevelopmental Cohort. Parental education and adverse experiences had unique associations with cortical surface area and subcortical volume as well as cognitive performance across several domains. Associations between parental education and several cognitive tasks were explained, in part, by variation in cortical surface area. In contrast, associations between adversity and cognitive tasks were explained primarily by variation in subcortical volume. A composite neurodevelopmental factor derived from principal component analysis of cortical thickness, cortical surface area, and subcortical volume mediated independent associations between both parental education and adverse experiences with reading, geometric reasoning, verbal reasoning, attention, and emotional differentiation tasks. Our analysis provides novel evidence that socioeconomic disadvantage and adversity influence neurodevelopmental pathways associated with cognitive outcomes through independent mechanisms.

## I. Introduction

Socioeconomic disadvantage is associated with adverse experiences, and yet disadvantage and adversity are distinct risk factors [1, 2] that both explain variation in brain development among youth [3, 4]. Disparities in childhood socioeconomic status (SES) related to parental education or household income have been associated with development of brain regions responsible for language [5, 6, 7, 8], executive function [9, 10, 11], memory [12, 13, 14, 8], and social cognition [15, 7, 16]. Studies have also found strong links between adverse experiences during childhood with structural and functional differences in the prefrontal cortex [17, 18, 19, 20, 21], which supports executive function, as well as the limbic system [15, 20, 22, 23, 24, 25, 26, 27, 28], which includes regions responsible for learning and socio-emotional processing.

SES and adverse experiences during childhood both also explain significant variation in cognitive outcomes across different domains [29, 30, 31, 32], which suggests that they both influence neural pathways related to cognitive function. Previous studies have observed that cortical surface area and white matter fiber tracts partially accounted for associations between family income and executive function [8, 11], while frontal and temporal lobe volume mediated associations between poverty and performance on academic achievement tests [33]. Studies of children who have been subjected to adversity have found that disparities in cerebellar volume explained links between early deprivation and memory [34], whereas reduced prefrontal cortical thickness mediated associations between institutionalization and attention disorders [35].

However, it is unclear whether links between SES and neurocognitive outcomes are entirely explained by adverse experiences, or other types of experiences, such as differences in cognitive stimulation or support from caregivers [1, 2, 16, 36]. Furthermore, the unique implications of distinct environmental exposures on neurodevelopmental factors that mediate cognitive outcomes are not well-understood.

Neuroanatomical development can be broadly classified according to cortical and subcortical structures. Gray matter volume in the cortex is a product of cortical thickness and surface area. Cortical surface area, cortical thickness, and subcortical volume are each characterized by different growth trajectories. Beginning in early childhood, cortical thickness decreases at varying rates across the brain until it stabilizes in early adulthood, likely due to synaptic pruning and myelination [37, 39]. Cortical surface area, on the other hand, expands from early childhood until middle adulthood when it begins contracting, likely, due to experience-driven synaptic pruning and outward pressure from myelination [40]. Maturation of subcortical volume is heterogeneous by region; for example, hippocampal and amygdala volumes increase until early adulthood before declining [41].

We examined neurodevelopmental pathways that mediated associations between parental education (PE), adverse experiences (AE), and cognitive outcomes in the Philadelphia Neurodevelopmental Cohort (PNC). The PNC collected genotypes and neuropsychological assessments, including an in-depth neurocognitive battery, from over 9,000 youth, 8 through 21 years of age, residing in the Philadelphia area between 2010 and 2014 [42, 43]. A subset of over 1,600 participants also underwent structural magnetic resonance imaging scans.

We first replicated a previous analysis correlating PE with cortical surface area and left hippocampal volume [8]. Next, we identified neuroimaging phenotypes for which PE and AE have significant unique effects, independent of age, sex, and ancestry. Then, we constructed mediation models [44] to assess the extent to which links between PE/AE and cognitive performance were accounted for by either cortical area or subcortical volume. Finally, we orthogonalized the cortical thickness, cortical surface area, and subcortical volume measures to characterize aspects of neurodevelopment associated with PE/AE.

The analysis sample consisted of 1, 413 PNC participants (see Table 1 for demographics) with data meeting availability and quality control criteria described in Materials and Methods (see Table S1 in **Supplementary Materials**). This sample is among the largest in the literature studying links between childhood experiences, brain structure, and cognitive outcomes. Moreover, our analysis is among a few in this field adjusting for genetic ancestry using genotype data to reduce bias from confounding associations between race, SES, and physiognomic measures such as brain structure [8, 11].

**Table 1:** PNC sample demographics.

|  | Full | History of adverse experiences |  |
| --- | --- | --- | --- |
|  |  | Yes | No |
| **Size** | 1413 | 766 (54.2%) | 647 (45.8%) |
| **Age (years)** |  |  |  |
| mean (sd) | 14.2 (3.5) | 14.9 (3.1) | 13.3 (3.8) |
| range | 8 to 21 | 8 to 21 | 8 to 21 |
| **Sex** |  |  |  |
| Male | 668 (47.3%) | 370 (48.3%) | 298 (46.1%) |
| Female | 745 (52.7%) | 396 (51.7%) | 349 (53.9%) |
| **Highest parental education (years)** |  |  |  |
| mean (sd) | 14.8 (2.6) | 14.5 (2.5) | 15.1 (2.6) |
| range | 8 to 20 | 8 to 20 | 9 to 20 |
| **Self-reported race** |  |  |  |
| European American | 639 (45.2%) | 300 (39.2%) | 339 (52.4%) |
| African American | 602 (42.6%) | 379 (49.5%) | 223 (34.5%) |
| Other/Unreported | 172 (12.2%) | 87 (11.4%) | 85 (13.1%) |

## II. Results

### Parental education and adverse experiences are associated with cortical area and gray matter volume

In our analysis, PE was defined as the maximum number of years of education completed by either parent. AE was defined as a binary variable indicating either a history of (AE=1) or no history of (AE=0) adverse experiences (i.e., fear of being harmed, being the victim of physical or sexual abuse, or witnessing violence or death; see Table S2 in **Supplementary Materials**). We adjusted for genetic ancestry using the first six principal components (PCs) of the genetic relatedness matrix in PNC’s genotype sample of 8,802 participants (see *Materials and Methods*). The first four genotype PCs corresponded to clustering with African, East Asian, South Asian, and mixed American ancestry groups, respectively, in the 1000 Genomes (phase 3) reference sample [45], while the fifth and sixth genotype PCs differentiated between subpopulations in the European and East Asian ancestry groups, respectively (see Figure S1 in **Supplementary Materials**).

Replicating a previous result in the Pediatric Imaging, Neurocognition, and Genetics (PING) cohort [8], we first observed a statistically significant positive relationship between PE and cortical surface area (see Table S3 in **Supplementary Materials**) in a linear regression model with covariates for age, sex, first six genotype PCs, PE, and age*×*PE (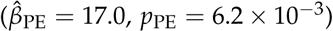). When regressing the same covariates against left hippocampal volume, which was significantly associated with PE in the PING cohort, the total effect from PE was significant (*p* = 0.035 under *F*-test of *H*_0_: *β*_PE_ = *β*_age × PE_ = 0) with a positive main effect (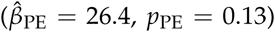). As in the previous work, PE did not have significant associations with cortical thickness, right hippocampal volume, and amygdala volumes in either hemisphere. Quadratic terms related to age were excluded as they did not contribute to unique variance in the PNC sample.

In our primary analysis of the whole brain (see Table 2), higher PE was associated with increased cortical area (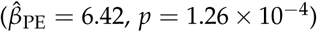) and increased subcortical volume 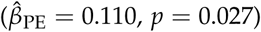, when adjusting for age, sex, first six genotype PCs, and AE. In contrast, the presence of AE was significantly associated with decreased cortical surface area 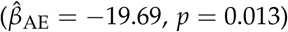 and subcortical volume 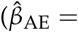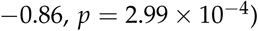, when adjusting for age, sex, first six genotype PCs, and PE. Although PE and AE are negatively correlated with each other (*r* = -0.115, *p* < 10 - 4), their significance in the same models suggests the two risk factors have independent effects on both measures. On the other hand, neither PE nor AE explained significant variation in mean cortical thickness. Including age × PE, age × AE, or PE × AE interactions did not significantly contribute to unique variation in the models for cortical thickness, cortical surface area, or subcortical volume.

**Table 2:** Associations between parental education and adverse experiences with cortical surface area, subcortical volume, and cortical thickness in the PNC (n=1,413).

| A. Cortical surface area (cm squared) |  |  |  |  |
| --- | --- | --- | --- | --- |
| | $\beta$ | $t$ | $p$ | partial $R^2$ |
| (Intercept) | 1851.17 | 63.13 | < 0.0001 | - |
| Age | -5.70 | -5.15 | < 0.0001 | 1.9% |
| SexF | -197.57 | -26.09 | < 0.0001 | 32.7% |
| PC1 | -61.38 | -13.40 | < 0.0001 | 11.3% |
| PC2 | -5.01 | -1.22 | 0.2219 | 0.1% |
| PC3 | 8.77 | 1.98 | 0.0478 | 0.3% |
| PC4 | 8.98 | 2.26 | 0.0242 | 0.4% |
| PC5 | 7.86 | 1.95 | 0.0515 | 0.3% |
| PC6 | 0.61 | 0.16 | 0.8711 | < 0.1% |
| PE | 6.42 | 3.84 | 0.0001 | 1.0% |
| AE | -19.69 | -2.50 | 0.0125 | 0.4% |

| B. Subcortical volume (cm cubed) |  |  |  |  |
| --- | --- | --- | --- | --- |
| | $\beta$ | $t$ | $p$ | partial $R^2$ |
| (Intercept) | 64.11 | 72.46 | < 0.0001 | - |
| Age | -0.17 | -5.00 | < 0.0001 | 1.8% |
| SexF | -4.27 | -18.71 | < 0.0001 | 20.0% |
| PC1 | -1.15 | -8.31 | < 0.0001 | 4.7% |
| PC2 | -0.10 | -0.84 | 0.4035 | < 0.1% |
| PC3 | 0.26 | 1.97 | 0.0492 | 0.3% |
| PC4 | 0.15 | 1.24 | 0.2143 | 0.1% |
| PC5 | 0.11 | 0.89 | 0.3720 | < 0.1% |
| PC6 | 0.24 | 2.12 | 0.0346 | 0.3% |
| PE | 0.11 | 2.20 | 0.0278 | 0.3% |
| AE | -0.86 | -3.63 | 0.0003 | 0.9% |

| C. Mean cortical thickness (mm) |  |  |  |  |
| --- | --- | --- | --- | --- |
| | $\beta$ | $t$ | $p$ | partial $R^2$ |
| (Intercept) | 2.86 | 145.92 | < 0.0001 | - |
| Age | -0.02 | -24.07 | < 0.0001 | 29.2% |
| SexF | 0.01 | 1.25 | 0.2104 | 0.1% |
| PC1 | -0.01 | -4.31 | < 0.0001 | 1.3% |
| PC2 | 0.00 | -1.81 | 0.0699 | 0.2% |
| PC3 | 0.00 | -0.14 | 0.8875 | < 0.1% |
| PC4 | 0.00 | 1.06 | 0.2906 | < 0.1% |
| PC5 | 0.00 | 1.11 | 0.2674 | < 0.1% |
| PC6 | 0.00 | 0.42 | 0.6726 | < 0.1% |
| PE | 0.00 | 1.20 | 0.2307 | 0.1% |
| AE | 0.00 | -0.65 | 0.5151 | < 0.1% |

SexF is a dummy variable equal to 1 for females and equal to 0 for males. PC1 through PC6 are first six PCs of genetic relatedness matrix over PNC's genotype sample. AE is a dummy variable equal to 1 for participants reporting a history of adverse experiences and equal to 0 otherwise. PE is highest parental education. $P$ -values are under $F$ -tests with 1 and 1,402 dof. Partial $R^2$ is percentage of variation in the model explained by an independent variable, after accounting for other covariates.

To identify regions of interest (ROIs) in the whole brain having strongest associations with PE/AE, we regressed the surface areas from 68 cortical ROIs and volumes from 14 subcortical ROIs against age, sex, first six genotype PCs, PE, and AE. The effects of PE and AE are not uniform across the brain (see Figure 1 and Table S4 in **Supplementary Materials**). Increased PE was associated with increased cortical ROI surface areas more so than subcortical ROI volumes. In contrast, the presence of AE was associated with decreases in volumes for more subcortical ROI and in surface areas of fewer cortical ROIs than increased PE. Several regions showed both a positive association with PE and a negative association with the presence of AE with p-values less than 0.05, after correcting for multiple testing with false discovery rate (FDR). These included the surface areas of the left inferior parietal and superior temporal gyri, the right superior temporal and caudal middle frontal gyri, as well as the volumes of the caudate in both hemispheres.

**Figure 1:**
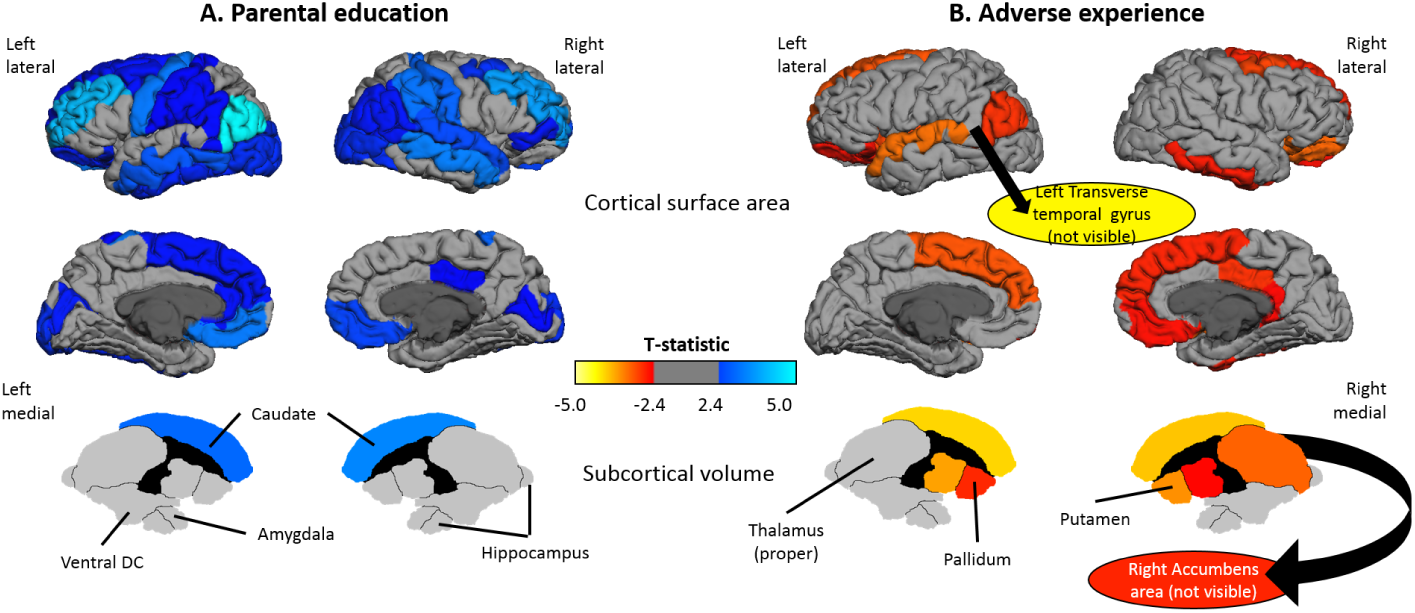
Regions associated with parental education or adverse experiences. Surface areas of 68 cortical ROIs and volumes of 14 subcortical were each regressed against age, sex, first six genotype PCs, PE, and AE. Tests were performed for the null hypotheses H_0_: β_PE_ = 0 and H_0_: β_AE_ = 0 in each model. Table S4 in **Supplementary Materials** reports summary statistics of associations significant at the 0.05 level, after correcting for 168 tests performed with FDR. (A) ROIs having significant association with parental education. (B) ROIs having significant association with adverse experiences.

### Correlations between cognitive performance with PE/AE risk factors and brain measures

We next examined associations between PE/AE and assessments across cognitive domains related to reading, complex cognition, executive function, and social cognition (see *Materials and Methods* for descriptions and Table S5 in **Supplementary Materials** for distributional statistics). PE was most strongly associated with performance on reading and reasoning tasks, explaining 17.3%, 6.8%, 15.0%, and 6.4% of unique variation (partial *R*^2^) on the WRAT, PMAT, PVRT, and PLOT, respectively, adjusting for age, sex, and AE (see Table 3). These observations are consistent with findings reporting that socioeconomic risk factors account for up to 20% of variation in overall cognitive performance [29]. In contrast, AE had a more muted association with cognitive performance — after accounting for age, sex, and PE, significant associations at the 0.05 level (unadjusted) were observed only in the WRAT and PMAT.

**Table 3:** Associations between parental education and adverse experiences with cognitive assessments in the PNC (n=1,413).

| Subdomain | Exam | n* | partial $R^2$ (sign($\beta$)) | |
| --- | --- | --- | --- | --- |
|  |  |  | PE | AE |
| Reading | WRAT<sup>a,b</sup> | 1380 | 17.3% (+) | 0.3% (–) |
| Geometric reasoning | PMAT<sup>a,c</sup> | 1371 | 6.8% (+) | 0.6% (–) |
| Verbal reasoning | PVRT<sup>a</sup> | 1369 | 15.0% (+) | 0.2% (–) |
| Spatial reasoning | PLOT<sup>a</sup> | 1252 | 6.4% (+) | <0.1% (+) |
| Working memory | LNB<sup>a</sup> | 1247 | 4.6% (+) | 0.2% (–) |
| Abstraction | PCET<sup>a</sup> | 1365 | 2.8% (+) | <0.1% (–) |
| Attention | PCPT<sup>a</sup> | 1254 | 2.4% (+) | <0.1% (+) |
| Spatial memory | VOLT<sup>a</sup> | 1350 | 3.0% (+) | <0.1% (+) |
| Face memory | PFMT<sup>a</sup> | 1324 | 1.8% (+) | <0.1% (+) |
| Emotional differentiation | PEDT<sup>a</sup> | 1357 | 2.7% (+) | <0.1% (–) |
| Emotional identification | PEIT | 1372 | 0.1% (+) | <0.1% (+) |
| Age differentiation | PADT<sup>a</sup> | 1370 | 2.2% (+) | 0.1% (+) |

Cognitive exam scores were regressed against age, sex, highest parental education (PE), and an indicator variable for a history of adverse experiences (AE). n\* is the number of participants in the neuroimaging sample (n=1,413) having valid scores on the respective exam. Partial $R^2$ is percentage of variation in the model explained by the respective independent variable, after accounting for other covariates. Significance of PE and AE in models of each cognitive exam was evaluated under *F*-tests with 1 and n\* – 5 dof. (a) Significance level of PE was less than $10^{-7}$ . (b) Significance level of AE was less than 0.05. (c) Significance level of AE was less than 0.01.

There were varying degrees of correlation between the cognitive assessments and cortical thickness, cortical area, or subcortical volume, after adjusting each cognitive and neuroanatomical measure for age, sex, and first six genotype PCs (see Figure S2 in **Supplementary Materials**). The highest correlations observed were between reading, geometric reasoning, and verbal reasoning (WRAT, PMAT, and PVRT) with either cortical surface area or subcortical volume. Cortical thickness was not significantly associated with any of the twelve cognitive exams.

### Brain measures mediate PE and AE effects on cognitive performance

We performed mediation analysis with either PE or AE as independent variables (IV), cortical area or subcortical volume as mediators, and scores on 12 cognitive exams as outcomes (see *Materials and Methods*). In the 48 respective pathways (2 IVs × 2 mediators × 12 outcomes), we estimated effects of the IV on the outcome through the mediator (indirect effect) and independent of the mediator (direct effect) through 100, 000 bootstrap simulations, while controlling for age, sex, first six genotype PCs, and AE (if PE was the IV) or PE (if AE was the IV). Averages, p-values, and 95% confidence intervals for the indirect effect, direct effect, and the ratio between the IE and the total effect in the bootstrap distributions for all 48 pathways are reported in Table S6 (**Supplementary Materials**).

Six pathways had an indirect effect that were significant at the 0.05 level, after correcting for multiple testing in 48 mediation models (Table 4 and Figure 2). Among them, cortical surface area mediated associations between PE and assessments on reading, geometric reasoning, verbal reasoning and working memory with indirect effects explaining 6.2%, 7.2%, 5.7%, and 7.2%, respectively, of the total effect. Subcortical volume significantly mediated the association between AE with reading and verbal reasoning. In these two pathways, AE did not have a significant direct effect on either assessment independent of age, sex, genetic ancestry, PE, and subcortical volume.

**Table 4:** Mediation pathways between parental education or adverse experiences, cortical surface area or subcortical volume, and cognitive outcomes.

| Independent variable | Mediator | Cognitive outcome |  | Estimate | *p* | 95% CI |
| --- | --- | --- | --- | --- | --- | --- |
| Parental education | Cortical surface area | Reading (WRAT) | Indirect effect | 0.56 | < 0.001 | 0.26 to 0.92 |
|  |  |  | Direct effect | 8.43 | < 0.001 | 6.88 to 9.99 |
|  |  |  | Indirect/Total effect | 0.062 | < 0.001 | 0.03 to 0.1 |
|  |  | Geometric reasoning (PMAT) | Indirect effect | 0.46 | < 0.001 | 0.18 to 0.81 |
|  |  |  | Direct effect | 5.88 | < 0.001 | 3.67 to 8.01 |
|  |  |  | Indirect/Total effect | 0.072 | < 0.001 | 0.03 to 0.14 |
|  |  | Verbal reasoning (PVRT) | Indirect effect | 0.49 | < 0.001 | 0.21 to 0.86 |
|  |  |  | Direct effect | 8.13 | < 0.001 | 6.35 to 9.93 |
|  |  |  | Indirect/Total effect | 0.057 | < 0.001 | 0.02 to 0.1 |
|  |  | Working memory (LNB) | Indirect effect | 0.39 | 0.003 | 0.11 to 0.77 |
|  |  |  | Direct effect | 4.98 | < 0.001 | 2.72 to 7.2 |
|  |  |  | Indirect/Total effect | 0.072 | 0.003 | 0.02 to 0.17 |
| Adverse experiences | Subcortical volume | Reading (WRAT) | Indirect effect | -2.33 | < 0.001 | -4.03 to -0.93 |
|  |  |  | Direct effect | 0.09 | 0.977 | -7.61 to 7.83 |
|  |  |  | Indirect/Total effect | 1.042 | 0.574 | -7.63 to 8.31 |
|  |  | Verbal reasoning (PVRT) | Indirect effect | -2.28 | < 0.001 | -3.96 to -0.91 |
|  |  |  | Direct effect | 1.68 | 0.681 | -6.65 to 10.34 |
|  |  |  | Indirect/Total effect | 3.815 | 0.912 | -8.25 to 8.26 |

**Figure 2:**
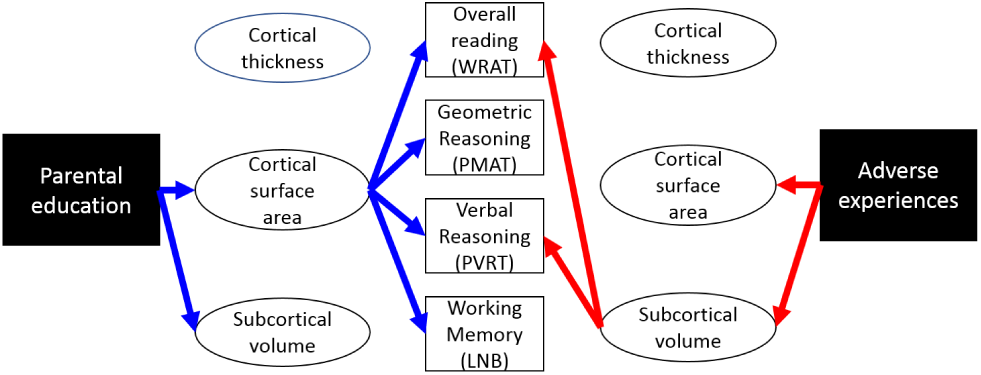
Pathways with significant indirect effects in mediation analysis.

### Mediation by orthogonal dimensions of neurodevelopment

Principal component analysis of the cortical thickness, cortical surface area, and subcortical volume measures, after residualizing each against age, sex, and first six genotype PCs, suggested three orthogonal neurodevelopmental (ND) factors related to expansion, contraction, and spatial differences (see Table 5 and Figure S3 in **Supplementary Materials**). While the expansion factor is roughly an average of cortical surface area and subcortical volume, the contraction factor primarily represents cortical thickness, which is subject to thinning during development. The spatial factor represents the difference between subcortical and cortical development. We termed the ND factors âĂŞ which are uncorrelated by design âĂŞ in this way due to the neurodevelopmental trajectories they represent.

**Table 5:** Loadings on neurodevelopmental (ND) factors through PCA of cortical thickness, cortical surface area, and subcortical volume.

| ND factor | Cortical thickness | Cortical surface area | Subcortical volume | % variation explained |
| --- | --- | --- | --- | --- |
| Expansion | -0.14 | 0.72 | 0.68 | 52.9% |
| Contraction | 0.92 | -0.15 | 0.35 | 36.1% |
| Spatial | -0.35 | -0.68 | 0.65 | 11.0% |

Prior to performing PCA, cortical thickness, cortical surface area, and subcortical volume were each respectively residualized against age, sex, and first six genotype PCs and then scaled to zero mean and unit variance.

We regressed the three ND factors against age, sex, the first six genotype PCs, PE, and AE (see Table 6). PE and AE were significantly associated with the expansion factor (*β*_PE_ = 0.05, *p*_PE_ = 0.0012;*β*_AE_ =*-*0.23,*p*_AE_ = 0.0009), whereas PE also had a significant association with the spatial factor (*β*_PE_ =*-*0.02, *p*_PE_ = 0.0055). Neither PE nor AE were associated with the contraction factor. These results were consistent with what we observed with the cortical thickness, cortical surface area, and subcortical volume measures that compose the ND factors (see Table 2).

**Table 6:** Associations between expansion, contraction, and spatial neurodevelopmental factor scores with parental education and adverse experiences.

A. Expansion factor
| | $\beta$ | $t$ | $p$ | partial $R^2$ |
| --- | --- | --- | --- | --- |
| (Intercept) | -0.69 | -2.66 | 0.0080 | - |
| Age | 0.01 | 0.81 | 0.4207 | < 0.1% |
| SexF | -0.02 | -0.22 | 0.8235 | < 0.1% |
| PC1 | 0.08 | 1.97 | 0.0487 | 0.3% |
| PC2 | -0.01 | -0.24 | 0.8105 | < 0.1% |
| PC3 | 0.01 | 0.19 | 0.8460 | < 0.1% |
| PC4 | -0.01 | -0.39 | 0.6991 | < 0.1% |
| PC5 | 0.01 | 0.41 | 0.6801 | < 0.1% |
| PC6 | -0.01 | -0.17 | 0.8633 | < 0.1% |
| PE | 0.05 | 3.25 | 0.0012 | 0.7% |
| AE | -0.23 | -3.31 | 0.0009 | 0.8% |

B. Contraction factor
| | $\beta$ | $t$ | $p$ | partial $R^2$ |
| --- | --- | --- | --- | --- |
| (Intercept) | -0.22 | -1.02 | 0.3058 | - |
| Age | 0.00 | 0.35 | 0.7264 | < 0.1% |
| sexF | -0.01 | -0.10 | 0.9237 | < 0.1% |
| PC1 | 0.03 | 0.79 | 0.4307 | < 0.1% |
| PC2 | 0.00 | -0.09 | 0.9267 | < 0.1% |
| PC3 | 0.00 | 0.08 | 0.9385 | < 0.1% |
| PC4 | 0.00 | -0.16 | 0.8755 | < 0.1% |
| PC5 | 0.01 | 0.17 | 0.8672 | < 0.1% |
| PC6 | 0.00 | -0.07 | 0.9444 | < 0.1% |
| PE | 0.02 | 1.26 | 0.2080 | 0.1% |
| AE | -0.08 | -1.44 | 0.1503 | 0.1% |

C. Spatial factor
| | $\beta$ | $t$ | $p$ | partial $R^2$ |
| --- | --- | --- | --- | --- |
| (Intercept) | 0.28 | 2.35 | 0.0187 | - |
| Age | 0.00 | 0.18 | 0.8590 | < 0.1% |
| SexF | 0.00 | -0.02 | 0.9805 | < 0.1% |
| PC1 | -0.02 | -1.13 | 0.2575 | < 0.1% |
| PC2 | 0.00 | 0.22 | 0.8231 | < 0.1% |
| PC3 | 0.00 | -0.12 | 0.9027 | < 0.1% |
| PC4 | 0.00 | 0.17 | 0.8625 | < 0.1% |
| PC5 | 0.00 | -0.18 | 0.8573 | < 0.1% |
| PC6 | 0.00 | 0.08 | 0.9373 | < 0.1% |
| PE | -0.02 | -2.78 | 0.0055 | 0.5% |
| AE | -0.02 | -0.73 | 0.4644 | < 0.1% |

Expansion, contraction, and spatial factors are derived from cortical thickness, cortical surface area, and subcortical volume through PCA. SexF is a dummy variable equal to 1 for females and equal to 0 for males. PC1-6 are first six PCs of genetic relatedness matrix over PNC's genotype sample. AE is a dummy variable equal to 1 for participants reporting an adverse experience and equal to 0 otherwise. PE is highest parental education. $P$ -values are under $F$ -tests with 1 and 1,402 dof. Partial $R^2$ is percentage of variation in the model explained by an independent variable, after accounting for other covariates.

We repeated the same mediation analysis performed above where cortical surface area and subcortical volumes were mediators in pathways linking PE/AE to cognitive exams, except using instead the expansion and spatial factors as mediators. Again, indirect effects were simulated for 48 mediation models through 100,000 bootstraps that include covariates for age, sex, genetic ancestry, and AE (if PE was the IV) or PE (if AE was the IV). Among the 48 models, there were 12 pathways with FDR-adjusted *p*-values less than 0.05 (see Figure 3 and Table S7 in **Supplementary Materials**) in which the expansion factor mediated associations between either PE or AE and the same six cognitive tasks related to reading, geometric reasoning, verbal reasoning, working memory, attention, and emotional differentiation. While direct effects of PE in these pathways were also significant, the direct effects of AE on the six exams were not significant, independent of age, sex, genetic ancestry, PE, and the expansion factor.

**Figure 3:**
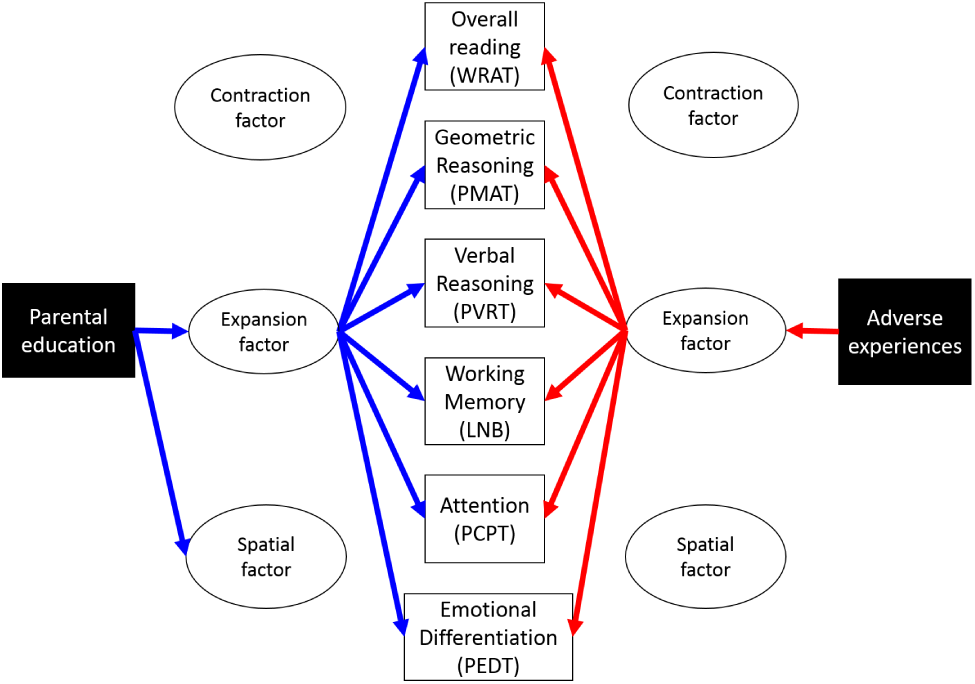
Pathways through neurodevelopmental factor scores having significant indirect effects in mediation analysis. See Table S7 in **Supplementary Materials** for summary statistics.

## III. Discussion

We first replicated a previous finding, in an independent sample, that PE is associated with variation in cortical surface area and left hippocampal volume [8]. This finding in the PNC sample further supports the observation that childhood SES factors play a critical role in neurodevelopment independent of age, sex, and genetic ancestry. We then explored similarities and differences in how various neurodevelopmental dimensions mediate the associations between PE and/or AE on cognitive performance. Although PE and

AE were negatively correlated (*r* = -0.12), our analysis targeting either risk factor included the other as a covariate to reduce bias in our inference.

Our main results confirm that lower parental education and previous adverse experiences are associated with worse cognitive performance primarily in domains related to reading and complex reasoning; these associations are partially accounted for by reduced cortical surface area and reduced subcortical volume. Moreover, PE and AE are distinct risk factors for neurodevelopment as PE is associated primarily with variation in cortical ROI surface area and AE is primarily associated with subcortical volumes. These observations suggest that there are spatial differences in the neurodevelopmental pathways that mediate associations between PE/AE and cognitive outcomes.

Statistically orthogonalizing cortical thickness, cortical area, and subcortical volume through principal components analysis allowed us to isolate commonalities and differences between the measures into neurodevelopmental factors. We found significant associations between PE/AE and the expansion factor, which represent a composite of cortical surface area and subcortical volume. This factor also significantly mediated relationships of both PE and AE with the same cognitive tasks. These observations suggest a common mechanism, related to differential rates in synaptic pruning and myelination during development, through which diminished PE and the presence of AE influence common aspects of neurocognitive development.

The mediation analysis suggested that variation in either cortical surface area or the expansion factor significantly explained between 4 and 8% of associations between PE and several cognitive exams. In contrast, links between AE and cognitive performances in reading and complex reasoning domains were driven primarily by variation in either subcortical volume or the expansion factor, when controlling for age, sex, genetic ancestry, and PE. One possible reason for the distinctive characteristics of these mediation pathways is that AE is not as pronounced a risk factor for diminished cognitive performance as is lower PE. However, it should be noted that our measure of PE was continuous, whereas our measure of AE was binary. It is therefore possible that we had less power to detect associations between AE and outcomes of interest. Nonetheless, these observations suggest that disparities in PE may have implications for neurocognitive development that are independent of adverse experiences.

Our results do not necessarily imply causal relationships between PE, AE, brain structure, and cognitive skills. First, PE is a distal marker for environmental factors such as nutrition, toxins, lack of caregiver support, learning impairments such as dyslexia or attention disorders that are linked to genetics, and fewer opportunities for cognitive enrichment that were not assessed in the present sample [1, 2]. Similarly, it is unlikely that (a) all types of AE have equal effects on neurocognitive development and (b) all individuals accurately report whether they have undergone an AE. Further investigations incorporating more complete descriptions of adverse experiences may provide greater insight into their influences on neurocognitive development in relation to SES.

Our study contributes to a growing literature on the role played by brain structure and physiology in explaining associations between environmental factors and cognition among youth [3, 46]. If disadvantage and adversity have unique implications for neurocognitive development, as our results suggest, then remediation is likely to be more effective at improving educational outcomes when targeted to specific risk profiles. Devising targeted early intervention strategies would help achieve the public policy goal of reducing effects of socioeconomic disparities on academic achievement among youth [47].

## IV. Methods and Materials

Raw neuroimaging scans, neuropsychological assessments, genotypes, and demographic information collected by the PNC study were obtained from dbGAP [43].

### Neuroimaging

T1-weighted structural scans were available for 1,598 participants, all of which were acquired with the same Siemens TIM Trio 3T scanner located at the Children’s Hospital of Philadelphia using a magnetization prepared, rapid-acquisition gradient-echo (MPRAGE) sequence (TR/TE =1810/3.5 milliseconds). Neuroimaging acquisition protocols and sample inclusion criteria have been previously described [42].

Structural scans were processed with the *recon-all* pipeline from FreeSurfer [48] (v5.3.0) to obtain morphometric measurements of the whole brain and regions of interest (ROIs) defined by the Desikan-Killiany atlas48. For quality control, we followed guidelines from the ENIGMA3 study (http://bit.ly/2q4FW4O) to identify 23 morphometric outliers and 83 participants with poor internal or external reconstructions who we excluded from further analysis. A scan was deemed to be a morphometric outlier if either total cortical surface area, mean cortical thickness, or subcortical volume was more than 2.698 standard deviations from the averages of the respective measures over scans for participants of the same age in the sample. This threshold is equivalent to two times the interquartile range for normal distributions. Authors CMM and JGM independently performed visual inspections of the internal and surface reconstructions. In general, reconstructions deemed to have inferior quality corresponded to scans exhibiting signs of excessive head motion.

### Cognitive assessments

Participants received a computerized battery of exams [43, 49] that included the Wide Range Achievement Test [50] (WRAT), Penn Matrix Reasoning Test [51] (PMAT), Penn Verbal Reasoning Test [52] (PVRT), and the Penn Line Orientation Test [51] (PLOT), all of which are related to complex cognition. Our analysis of the WRAT used raw scores on the reading subscale of the WRAT-IV, which tested word reading and sentence comprehension skills. PMAT and PVRT scores corresponded to the number of correct responses to analogy problems related, respectively, to geometric reasoning and verbal skills. In the PLOT, test-takers rotated a pair of lines until their angle matched a target image.

The Letter *N*-back Test [53] (LNB), Penn Conditional Exclusion Test [54] (PCET), and the Penn Continuous Performance Test [51, 55, 56] (PCPT) assessed working memory, abstraction, and attention, respectively. In the LNB, participants were presented with sequences of uppercase letters with a stimulus duration of 0.5 seconds in trials separated by 2.5 seconds. The LNB score was the total number of correct responses during the 1-back and 2-back conditions, where participants responded if the letter was identical to the letter presented in the previous trial or two trials previously. The PCET required participants use one of three geometric sorting principles not made explicit to them. Participants were assigned an accuracy score based on performance on the PCET. The PCPT measures attention through a rapid presentation of sixty displays consisting of seven segments for one-second durations. In the first half of the test, participants pressed the space bar if they saw a number; in the second half of the test, participants pushed the space bar if they saw a letter.

The Visual Object Learning Test [57] (VOLT), Penn Word Memory Test [58] (PWMT), and Penn Face Memory Test [58] (PFMT) assessed spatial, verbal and face memory, respectively. In the VOLT, participants were asked to memorize 20 targets consisting of Euclidean shapes and then identify the targets from a set of images that included 20 foils. A participant’s score was the number of correctly identified targets and correctly rejected foils. The PWMT and PFMT have the same testing and scoring procedures except that the targets and foils in the respective exams were words and images of faces instead of shapes.

The Penn Emotion Identification Test (PEIT), Penn Emotion Differentiation Test (PEDT), and the Penn Age Differentiation Test (PADT) were used to measure social cognition [59, 60, 61]. Participants were presented with pictures of faces with happy, sad, angry, fearful, or neutral expressions in the PEIT and the PEDT. In the PEIT, participants identified emotions on faces, which were presented one at a time. In the PEDT, participants identified which of two simultaneously presented faces had greater emotional intensity or indicated the intensities were equivalent. In the PADT, participants identified the older of two simultaneously presented faces or indicated the two faces were equivalent in age.

Only scores from completed exams were included in analyses. Raw scores were divided by the maximum possible on the given form and then standardized to zero mean and standard deviation 100. Standardized scores more than three standard deviations from the mean were excluded from analysis. Because the PNC study administered multiple forms of the PMAT, PVRT, PLOT, PEDT, PEIT, and PADT, standardization was performed separately over each form for those exams. Table S5 in the **Supplementary Materials** provides test descriptions, the number of participants with valid scores available for analysis, and sample statistics for mean, standard deviation, skewness and excess kurtosis. After inspecting the distribution of the scaled scores, and after correcting for age, sex, and genetic ancestry, we excluded the PWMT from further analysis after finding that its scores severely violated normality assumptions required for our statistical analyses.

### Genetic ancestry

PNC participants were genotyped with several platforms, including Illumina’s HumanHap550, 610- Quad, and OmniExpress, as well as Affymetrix’s Axiom and 6.0 chips (see Table S8 in **Supplementary Materials** for summary). The genotype collection pipeline has been described previously [62, 63] and further details are available at www.caglab.org. Genotypes from each platform were separately imputed to a reference panel of *n* = 2, 504 participants in the 1000 Genomes Phase 3 (1KG) project [45] with *genipe* [64] After applying quality control filters and performing imputation (see **Supplementary Materials** for details), we combined data for 376, 373 single nucleotide polymorphisms, having minor allele frequencies greater than 0.05 in imputed datasets, for 8, 802 PNC participants as well as the 2, 504 individuals in the 1KG reference sample. We performed PCA on the combined dataset, after pruning to 31, 930 SNPs with low linkage disequilibrium (*r*^2^ *<* 0.1), using PLINK v1.9 [65]. Violin plots illustrate that the PC scores clustered with self-reported races in the PNC and ancestry groups in the 1KG reference sample (see Figure S1 in **Supplementary Materials**).

### Statistical analysis

We fit linear regression models of total subcortical gray matter volume, total cortical surface area, and mean cortical thickness to age, sex, the first six PCs of genotype data, PE, and AE over the PNC neuroimaging sample (*n* = 1, 413). Covariates’ significance in each model was assessed using a *F*-test with 1 and 1, 402 degrees of freedom (dof). We also regressed surface areas of 68 cortical ROIs and volumes of 14 subcortical ROIs against the same covariates. Tests were performed for the null hypotheses *H*_0_: *β*_PE_ = 0 and *H*_0_: *β*_AE_ = 0 in a total of 84 (66 + 16) models of ROIs measures. To correct for multiple testing, observed *p*-values in the 168 (2 × 84) tests performed in this analysis (*F*-test with 1 and 1, 402 dof) were adjusted using false discovery rate.

We performed mediation analysis to detect whether structural brain measures (mediator) explained associations between either PE or AE (independent variable [IV]) on cognitive exam scores (outcome). In the framework that we used, models of a mediator and an outcome were fit separately over the sample. The mediator was regressed against the IV and nuisance covariates, and outcome was regressed against the mediator, IV, and nuisance covariates. Let *Û*_*i*_:= *f* (*X*_*i*_, *W*_*i*_) and *Ŷ* _*i*_:= *g*(*X*_*i*_, *U*_*i*_, *W*_*i*_) denote predictions from the mediator and outcome models, respectively, where *X*_*i*_ is the independent variable, *U*_*i*_ is the mediating variable, and *W*_*i*_ stands for nuisance covariates. We estimated the indirect effect (IE) and direct effect (DE), which are defined as

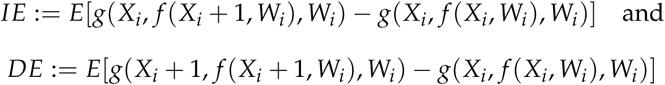

respectively, through 100,000 non-parametric bootstraps with the R-package *mediation* [66] in a total of 48 mediation models with either PE or AE as the IV, cortical surface area or subcortical volume as mediators, and scores on 12 cognitive exams as outcomes. Covariates in both the outcome and mediator models were age, sex, the first six genotype PCs, PE, and AE.

We used PCA to statistically orthogonalize cortical thickness, cortical area, and subcortical volume after each measure was first residualized against age, sex, and the first six genotype PCs with linear regression and then scaled to zero mean and unit variance. Scores for three orthogonal neurodevelopmental factors (ND) for each participant i were

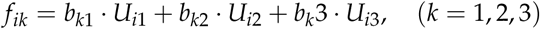

*where U*_*i*1_, *U*_*i*2_, and *U*_*i*3_ represent the three residualized brain measures and *b*_*k*1_, *b*_*k*2_, and *b*_*k*3_ are estimated PC loadings on components *k* = 1, 2, 3, respectively. We then repeated our analysis using three ND scores (*f*_*ik*_), instead of cortical thickness, cortical area, and subcortical volume. All analyses were performed in R unless otherwise noted. Scripts are available by request.

## V. Supplementary information

### Genotype analysis

dbGAP provided genotypes for 9, 002 PNC participants distributed over thirteen datasets (see Table S8 for summary). Major and minor allele designations for the Illumina platforms were converted from A/B format, as given on dbGAP, to A/T/C/G format using strand information for the respective platforms obtained from http://www.well.ox.ac.uk/wrayner/strand/. SNP positions on all platforms were updated to HG build 19 with the LiftOver script in Python obtained from http://genome.sph.umich.edu/wiki/LiftOver. SNPs with minor allele frequency, Hardy-Weinberg equilibrium *p*-value, and genotyping call rate less than 0.05, 10*-*5, and 0.05, respectively, were filtered from three largest datasets. Samples were also filtered so the minimum genotype call rate was 95%, among whom 32 had neuroimaging scans available.

Genotypes were then imputed to approximately 13.7 million SNPs across the genome from 2, 504 participants in the publicly available 1KG reference sample using genipe [64]. Genotypes from the imputed PNC datasets (*n* = 8, 802) and 1KG project (*n* = 2, 504) were combined into a single dataset while restricting to 376,373 SNPs having minor allele frequency greater than 0.05 on all ten imputed datasets and were also directly genotyped by either the HumanHap550-v3, 610-Quad, and OmniExpress platforms, which were used for 83% of participants in the full sample. PCA was performed on the combined dataset, after pruning to 31, 930 SNPs with low linkage disequilibrium (*r*^2^ *<* 0.1).

**Table S1.**
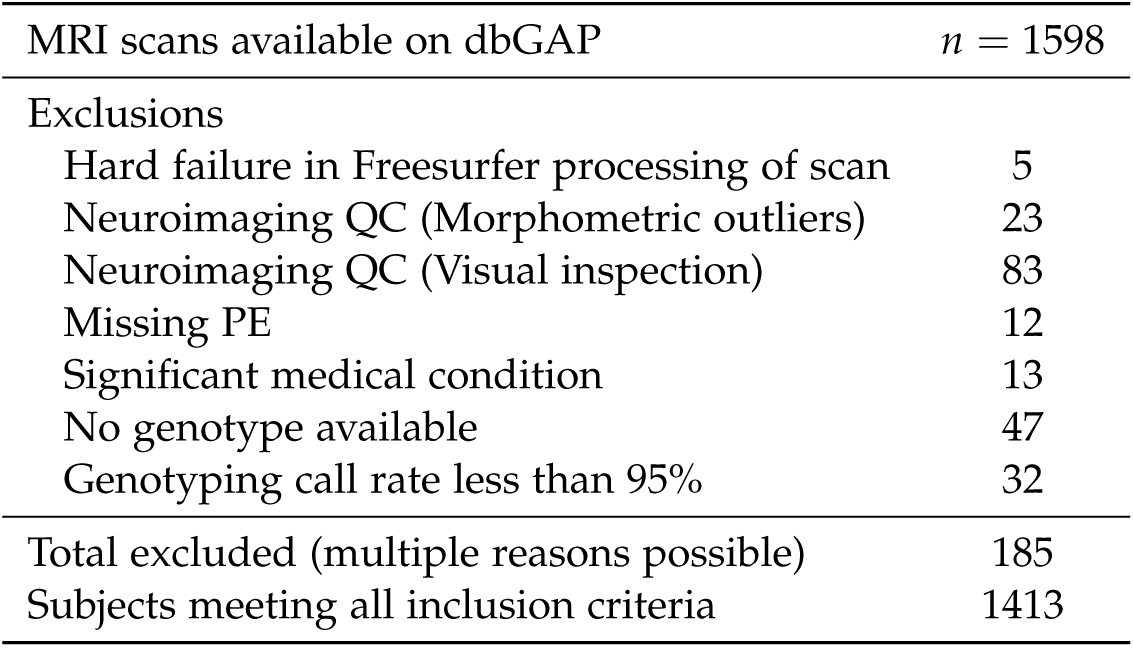
PNC sample inclusion criteria.

| MRI scans available on dbGAP | *n* = 1598 |
| --- | --- |
| Exclusions |  |
| Hard failure in Freesurfer processing of scan | 5 |
| Neuroimaging QC (Morphometric outliers) | 23 |
| Neuroimaging QC (Visual inspection) | 83 |
| Missing PE | 12 |
| Significant medical condition | 13 |
| No genotype available | 47 |
| Genotyping call rate less than 95% | 32 |
| Total excluded (multiple reasons possible) | 185 |
| Subjects meeting all inclusion criteria | 1413 |

**Table S2.** Breakdown of adverse experiences in the PNC (n=1,413).

| Question ID | Adverse event | # citing this event |
| --- | --- | --- |
| PTD001 | Fear of harm due to natural disaster | 45 |
| PTD002 | Fear of harm towards self or someone close (e.g. family violence) | 177 |
| PTD003 | Being attacked or badly beaten | 97 |
| PTD004 | Upset at forced sexual encounter | 49 |
| PTD005 | Attacked sexually or raped | 38 |
| PTD006 | Threatened with a weapon | 90 |
| PTD007 | Been in bad accident | 156 |
| PTD008 | Seen/heard someone get killed or hurt badly(outside of movies) | 303 |
| PTD009 | Upset at seeing dead body of someone close | 347 |
| PTD020* | Provided age of experiencing traumatic event but not specific type | 86 |

Subjects may have reported multiple types of adverse or traumatic experiences. \*Subjects reported age of experiencing of an adverse event but did not indicate its type.

**Table S3.** Replication of Noble, et al. (2015) in the PNC (n=1,413).

A. Model of cortical area (cm squared)
| | $\beta$ | $t$ | $p$ | partial $R^2$ |
| --- | --- | --- | --- | --- |
| (Intercept) | 1693.37 | 18.25 | < 0.0001 | - |
| Age | 4.33 | 0.70 | 0.4862 | 2.4% |
| SexF | -196.43 | -25.96 | < 0.0001 | 32.5% |
| PC1 | -62.81 | -13.83 | < 0.0001 | 12.0% |
| PC2 | -5.47 | -1.33 | 0.1841 | 0.1% |
| PC3 | 8.42 | 1.90 | 0.0578 | 0.3% |
| PC4 | 9.23 | 2.32 | 0.0206 | 0.4% |
| PC5 | 6.91 | 1.71 | 0.0873 | 0.2% |
| PC6 | 0.64 | 0.17 | 0.8655 | < 0.1% |
| PE | 17.00 | 2.74 | 0.0062 | 1.1% |
| age × PE | -0.73 | -1.75 | 0.0805 | 0.2% |

B. Model of left hippocampal volume (mm cubed)
| | $\beta$ | $t$ | $p$ | partial $R^2$ |
| --- | --- | --- | --- | --- |
| (Intercept) | 3836.91 | 14.71 | < 0.0001 | - |
| Age | 15.84 | 0.91 | 0.3648 | < 0.1% |
| SexF | -272.16 | -12.79 | < 0.0001 | 10.4% |
| PC1 | -121.41 | -9.50 | < 0.0001 | 6.1% |
| PC2 | -26.95 | -2.33 | 0.0199 | 0.4% |
| PC3 | 8.79 | 0.70 | 0.4810 | < 0.1% |
| PC4 | 8.60 | 0.77 | 0.4428 | < 0.1% |
| PC5 | 9.89 | 0.87 | 0.3842 | < 0.1% |
| PC6 | 10.26 | 0.96 | 0.3355 | < 0.1% |
| PE | 26.42 | 1.52 | 0.1297 | 0.4% |
| age × PE | -1.04 | -0.89 | 0.3723 | < 0.1% |

Test of total PE effect: F-stat=3.37, p-value=0.03481
SexF is a dummy variable equal to 1 for females and equal to 0 for males. PC1-6 are the first six PCs of the genetic relatedness matrix in PNC's genotype sample. AE is a dummy variable equal to 1 for participants reporting an adverse experience and equal to 0 otherwise. PE is highest parental education. F-tests with 1 and 1,402 dof were performed to evaluate significance ( $p$ ) for covariates in each model. F-tests with 2 and 1,402 dof were performed for the null hypothesis $H_0 : \beta_{PE} = \beta_{age \times PE} = 0$ in each model. Partial $R^2$ is percentage of variation in the model explained by an independent variable, after accounting for other covariates.

**Table S4.**
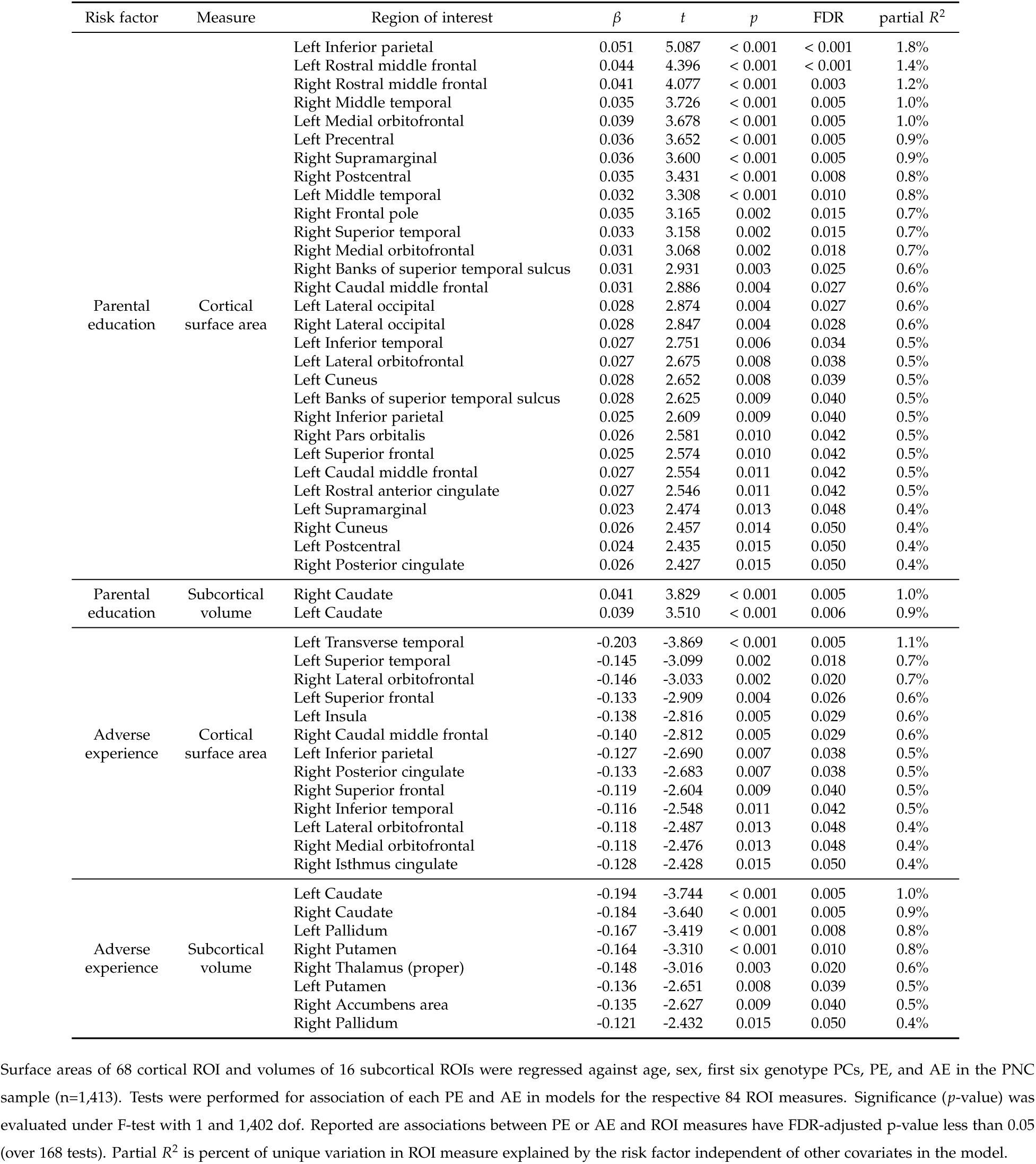
Associations between cortical ROI surface areas and subcortical ROI volumes with parental education and adverse experiences in the PNC (n=1,413).

**Table S5.** Distribution of cognitive assessments in the PNC (n=1,413).

| Domain | Exam | Subdomain | Full name | # forms | # questions* | Summary statistics of scores in sample |  |  |
| --- | --- | --- | --- | --- | --- | --- | --- | --- |
|  |  |  |  |  |  | n* | Skew | Excess kurt. |
| Complex cognition | WRAT | Reading | Wide Range Achievement Test | 1 | 70 | 1380 | -0.45 | -0.36 |
|  | PMAT | Geometric reasoning | Penn Verbal Reasoning Test | 4 | 18 or 24 | 1371 | 0.20 | -0.42 |
|  | PVRT | Verbal reasoning | Penn Matrix Reasoning Test | 4 | 15 | 1369 | -0.60 | -0.48 |
|  | PLOT | Spatial reasoning | Penn Line Orientation Test | 2 | 12 or 24 | 1252 | -0.02 | -0.65 |
| Executive function | LNB | Working memory | Letter n-back test | 1 | 20 | 1247 | -1.06 | 0.65 |
|  | PCET | Abstraction | Penn Conditional Exclusion Test | 1 | * | 1365 | -0.58 | -0.23 |
|  | PCPT | Attention | Penn Continuous Performance Test | 1 | 60 | 1254 | -1.23 | 1.14 |
| Episodic memory | VOLT | Spatial memory | Visual object learning test | 1 | 20 | 1350 | -0.22 | -0.58 |
|  | PWMT | Word memory | Penn Word memory Test | 1 | 40 | 1342 | -1.05 | 0.70 |
|  | PFMT | Face memory | Penn Face memory Test | 1 | 40 | 1324 | -0.16 | -0.48 |
| Social cognition | PEDT | Emotional differentiation | Penn Emotional Differentiation Test | 2 | 36 or 60 | 1357 | -0.48 | -0.11 |
|  | PEIT | Emotional identification | Penn Emotional Identification Test | 4 | 40 | 1372 | -0.65 | 0.28 |
|  | PADT | Age differentiation | Penn Age Differentiation Test | 2 | 36 or 60 | 1370 | -0.51 | 0.07 |

Only scores on exams flagged as being valid were included in the analysis. n\* is the number of participants with valid scores in the respective assessment. Raw scores were the number of correctly answered questions on all assessments except the PCET, for which an accuracy score was reported. PNC administered multiple forms of the PMAT, PVRT, PLOT, PEDT, PEIT, and PADT. Forms for the PMAT, PLOT, PEDT, and PADT had different numbers of questions. Scores were divided by the maximum possible on the given form and then standardized to zero mean and standard deviation 100. Standardized scores more than three standard deviations from the mean were excluded from the analysis. \*A normalized accuracy score was reported for the PCET.

**Table S6.**
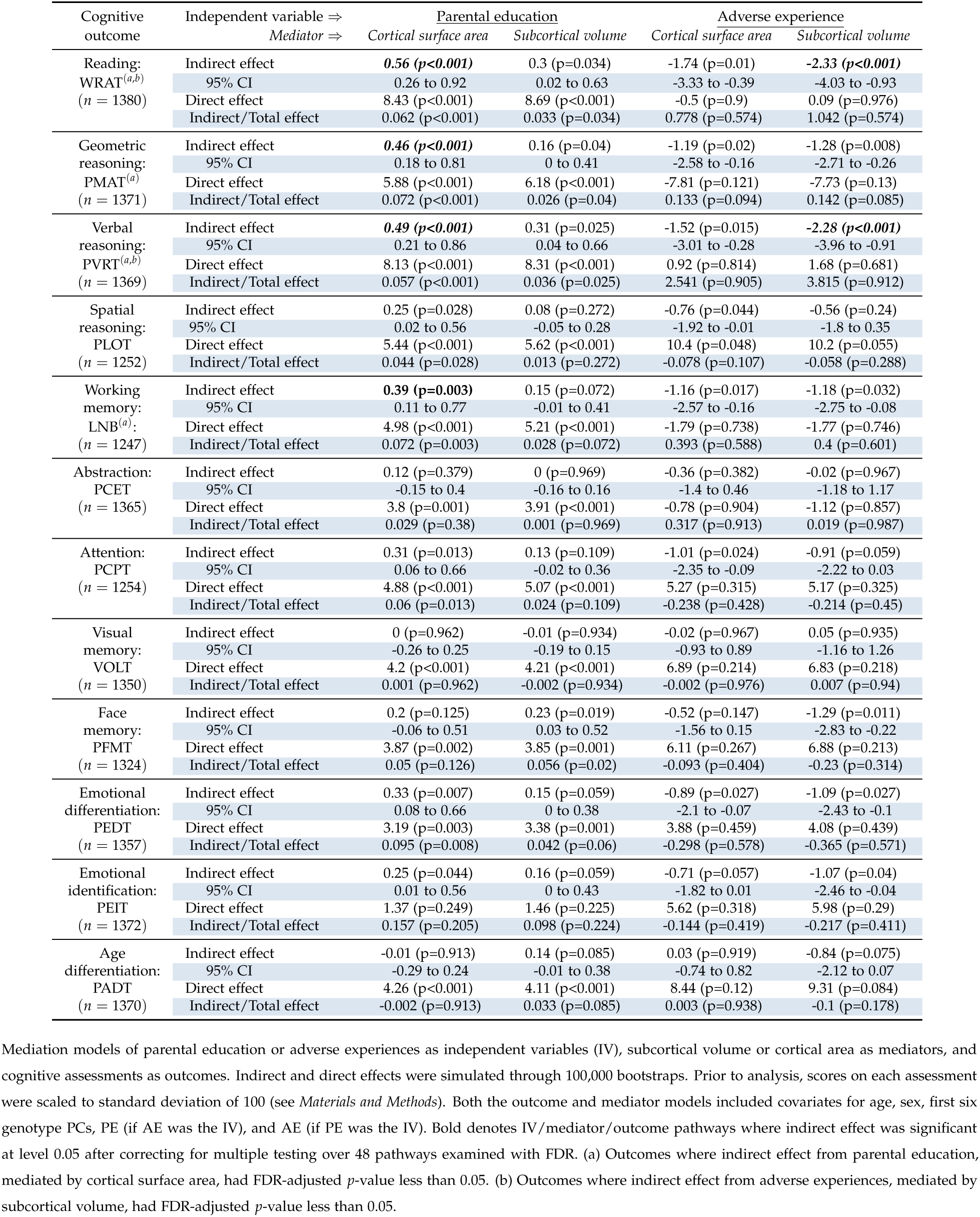
Mediation analysis for parental education/adverse experiences, cortical surface area/subcortical volume, and cognitive outcomes in the PNC (n=1,413).

**Table S7.** Mediation analysis for parental education/adverse experiences, expansion/spatial ND factors, and cognitive outcomes.

| Cognitive outcome | Independent variable ⇒<br>Mediator ⇒ | Parental education |  | Adverse experience |  |
| --- | --- | --- | --- | --- | --- |
|  |  | Expansion factor | Spatial factor | Expansion factor | Spatial factor |
| Reading:<br>WRAT<sup>(a)</sup><br>(*n* = 1380) | Indirect effect | **0.51 (*p*<0.001)** | 0.06 (*p*=0.387) | **-2.44 (*p*<0.001)** | 0.06 (*p*=0.706) |
|  | 95% CI | 0.19 to 0.88 | -0.07 to 0.21 | -4.26 to -0.92 | -0.21 to 0.45 |
|  | Direct effect | 8.48 (*p*<0.001) | 8.93 (*p*<0.001) | 0.2 (*p*=0.964) | -2.29 (*p*=0.558) |
|  | Indirect/Total effect | 0.057 (*p*<0.001) | 0.006 (*p*=0.387) | 1.092 (*p*=0.566) | -0.026 (*p*=0.87) |
| Geometric reasoning:<br>PMAT<sup>(a)</sup><br>(*n* = 1371) | Indirect effect | **0.36 (*p*<0.001)** | 0.1 (*p*=0.29) | **-1.57 (*p*=0.002)** | 0.11 (*p*=0.589) |
|  | 95% CI | 0.11 to 0.69 | -0.09 to 0.31 | -3.11 to -0.45 | -0.26 to 0.67 |
|  | Direct effect | 5.98 (*p*<0.001) | 6.24 (*p*<0.001) | -7.43 (*p*=0.142) | -9.12 (*p*=0.074) |
|  | Indirect/Total effect | 0.057 (*p*<0.001) | 0.016 (*p*=0.29) | 0.175 (*p*=0.077) | -0.013 (*p*=0.624) |
| Verbal reasoning:<br>PVRT<sup>(a)</sup><br>(*n* = 1369) | Indirect effect | **0.49 (*p*=0.001)** | -0.01 (*p*=0.897) | **-2.27 (*p*=0.002)** | -0.01 (*p*=0.968) |
|  | 95% CI | 0.17 to 0.88 | -0.16 to 0.14 | -3.99 to -0.82 | -0.32 to 0.3 |
|  | Direct effect | 8.13 (*p*<0.001) | 8.63 (*p*<0.001) | 1.67 (*p*=0.687) | -0.59 (*p*=0.915) |
|  | Indirect/Total effect | 0.056 (*p*<0.001) | -0.001 (*p*=0.897) | 3.801 (*p*=0.905) | 0.016 (*p*=0.993) |
| Spatial reasoning:<br>PLOT<br>(*n* = 1252) | Indirect effect | 0.18 (*p*=0.067) | 0.14 (*p*=0.142) | -0.81 (*p*=0.069) | 0.11 (*p*=0.67) |
|  | 95% CI | -0.01 to 0.45 | -0.04 to 0.39 | -2.09 to 0.05 | -0.39 to 0.78 |
|  | Direct effect | 5.52 (*p*<0.001) | 5.56 (*p*<0.001) | 10.45 (*p*=0.049) | 9.53 (*p*=0.07) |
|  | Indirect/Total effect | 0.031 (*p*=0.067) | 0.024 (*p*=0.142) | -0.084 (*p*=0.131) | 0.011 (*p*=0.69) |
| Working memory:<br>LNB<sup>(a)</sup><br>(*n* = 1247) | Indirect effect | **0.33 (*p*=0.004)** | 0.02 (*p*=0.793) | **-1.54 (*p*=0.005)** | 0.03 (*p*=0.859) |
|  | 95% CI | 0.08 to 0.69 | -0.17 to 0.21 | -3.21 to -0.35 | -0.32 to 0.5 |
|  | Direct effect | 5.03 (*p*<0.001) | 5.34 (*p*<0.001) | -1.41 (*p*=0.796) | -2.98 (*p*=0.583) |
|  | Indirect/Total effect | 0.061 (*p*=0.004) | 0.004 (*p*=0.793) | 0.522 (*p*=0.587) | -0.009 (*p*=0.936) |
| Abstraction:<br>PCET<br>(*n* = 1365) | Indirect effect | 0.07 (*p*=0.533) | 0.04 (*p*=0.652) | -0.32 (*p*=0.538) | 0.02 (*p*=0.888) |
|  | 95% CI | -0.16 to 0.31 | -0.15 to 0.25 | -1.5 to 0.74 | -0.31 to 0.46 |
|  | Direct effect | 3.84 (*p*<0.001) | 3.87 (*p*<0.001) | -0.82 (*p*=0.9) | -1.17 (*p*=0.85) |
|  | Indirect/Total effect | 0.017 (*p*=0.533) | 0.011 (*p*=0.653) | 0.281 (*p*=0.946) | -0.021 (*p*=0.992) |
| Attention:<br>PCPT<sup>(a)</sup><br>(*n* = 1254) | Indirect effect | **0.28 (*p*=0.011)** | -0.04 (*p*=0.671) | **-1.27 (*p*=0.011)** | 0.02 (*p*=0.897) |
|  | 95% CI | 0.05 to 0.61 | -0.24 to 0.15 | -2.74 to -0.21 | -0.3 to 0.44 |
|  | Direct effect | 4.91 (*p*<0.001) | 5.23 (*p*<0.001) | 5.53 (*p*=0.288) | 4.24 (*p*=0.413) |
|  | Indirect/Total effect | 0.054 (*p*=0.011) | -0.007 (*p*=0.671) | -0.299 (*p*=0.418) | 0.004 (*p*=0.944) |
| Visual memory:<br>VOLT<br>(*n* = 1350) | Indirect effect | -0.02 (*p*=0.862) | 0.08 (*p*=0.343) | 0.09 (*p*=0.868) | 0.06 (*p*=0.785) |
|  | 95% CI | -0.26 to 0.19 | -0.09 to 0.3 | -1.03 to 1.23 | -0.36 to 0.61 |
|  | Direct effect | 4.23 (*p*<0.001) | 4.12 (*p*<0.001) | 6.78 (*p*=0.223) | 6.81 (*p*=0.224) |
|  | Indirect/Total effect | -0.004 (*p*=0.862) | 0.02 (*p*=0.343) | 0.014 (*p*=0.892) | 0.009 (*p*=0.819) |
| Face memory:<br>PFMT<br>(*n* = 1324) | Indirect effect | 0.28 (*p*=0.019) | -0.11 (*p*=0.182) | -1.04 (*p*=0.023) | -0.09 (*p*=0.722) |
|  | 95% CI | 0.04 to 0.6 | -0.34 to 0.04 | -2.43 to -0.1 | -0.73 to 0.41 |
|  | Direct effect | 3.79 (*p*=0.002) | 4.18 (*p*<0.001) | 6.64 (*p*=0.23) | 5.68 (*p*=0.298) |
|  | Indirect/Total effect | 0.069 (*p*=0.02) | -0.027 (*p*=0.183) | -0.186 (*p*=0.323) | -0.016 (*p*=0.82) |
| Emotional differentiation:<br>PEDT<sup>(a)</sup><br>(*n* = 1357) | Indirect effect | **0.29 (*p*=0.007)** | 0.03 (*p*=0.738) | **-1.23 (*p*=0.008)** | 0.03 (*p*=0.888) |
|  | 95% CI | 0.07 to 0.59 | -0.16 to 0.23 | -2.6 to -0.23 | -0.32 to 0.45 |
|  | Direct effect | 3.23 (*p*=0.003) | 3.49 (*p*=0.001) | 4.22 (*p*=0.419) | 2.96 (*p*=0.574) |
|  | Indirect/Total effect | 0.082 (*p*=0.008) | 0.008 (*p*=0.739) | -0.411 (*p*=0.563) | 0.009 (*p*=0.976) |
| Emotional identification:<br>PEIT<br>(*n* = 1372) | Indirect effect | 0.23 (*p*=0.04) | 0.09 (*p*=0.328) | -0.97 (*p*=0.041) | 0.1 (*p*=0.644) |
|  | 95% CI | 0.01 to 0.54 | -0.08 to 0.31 | -2.29 to -0.03 | -0.27 to 0.72 |
|  | Direct effect | 1.39 (*p*=0.245) | 1.54 (*p*=0.203) | 5.89 (*p*=0.296) | 4.81 (*p*=0.391) |
|  | Indirect/Total effect | 0.142 (*p*=0.207) | 0.053 (*p*=0.442) | -0.199 (*p*=0.41) | 0.02 (*p*=0.765) |
| Age differentiation:<br>PADT<br>(*n* = 1370) | Indirect effect | 0.09 (*p*=0.428) | -0.12 (*p*=0.152) | -0.36 (*p*=0.431) | -0.09 (*p*=0.705) |
|  | 95% CI | -0.14 to 0.34 | -0.34 to 0.04 | -1.41 to 0.52 | -0.74 to 0.39 |
|  | Direct effect | 4.16 (*p*<0.001) | 4.37 (*p*<0.001) | 8.82 (*p*=0.103) | 8.56 (*p*=0.109) |
|  | Indirect/Total effect | 0.021 (*p*=0.428) | -0.027 (*p*=0.152) | -0.042 (*p*=0.484) | -0.011 (*p*=0.747) |

Mediation models of parental education or adverse experiences as independent variables (IV), expansion or spatial ND factors as mediators, and cognitive assessments as outcomes. Indirect and direct effects were simulated through 100,000 bootstraps. Prior to analysis, scores on each assessment were scaled to standard deviation of 100 (see *Materials and Methods*). Both the outcome and mediator models included covariates for age, sex, first six genotype PCs, PE (if AE was the IV), and AE (if PE was the IV). Bold denotes IV/mediator/outcome pathways where indirect effect was significant at level 0.05 after correcting for multiple testing over 48 pathways examined with FDR. (a) Outcomes where indirect effect from either parental education or adverse experiences, mediated by the expansion ND factor, had FDR-adjusted *p*-value less than 0.05.

**Table S8.** Summary of PNC’s genotype sample (n=9,002).

| Dataset | Array | # SNPs | Available from dbGAP |  | After QC<sup>b</sup> |  |
| --- | --- | --- | --- | --- | --- | --- |
|  |  |  | # participants |  | # in sample for PCA |  |
|  |  |  | Full | Imaging<sup>a</sup> | Full | Imaging<sup>a</sup> |
| GO-Quad | Human610-Quadv1-B (Illumina) | 620901 | 3802 | 693 | 3734 | 684 |
| GO-v3 | HumanHap550v3-A (Illumina) | 561466 | 1913 | 325 | 1862 | 302 |
| GO-Omni | HumanOmniExpress-12v1-A (Illumina) | 733202 | 1657 | 293 | 1653 | 293 |
| Axiom | Affymetrix Axiom | 567096 | 722 | 114 | 711 | 114 |
| GO-v1 | HUMANHAP550-11218540-C (Illumina) | 555352 | 555 | 84 | 547 | 84 |
| GO-1Mduo | Human1M-Duov3-B (Illumina) | 1199187 | 141 | 16 | 141 | 16 |
| Affy6.0 | Affymetrix 6.0 | 909622 | 66 | 15 | 66 | 15 |
| GO-Quad-set2 | Human610-Quadv1-B (Illumina) | 620901 | 40 | 5 | 40 | 5 |
| GO-v3-set2 | HumanHap550v3-A (Illumina) | 561466 | 31 | 3 | 31 | 3 |
| GO-v1-set2 | HUMANHAP550-11218540-C (Illumina) | 555352 | 17 | 3 | 17 | 3 |
| GO-Omni-set2 | HumanOmniExpress-12v1-A (Illumina) | 733202 | 37 | 0 | - | - |
| GO-OMNI-12v1-1 | HumanOmniExpress-12v1-1-B (Illumina) | 719665 | 18 | 0 | - | - |
| GO-OEE | HumanOmniExpressExome-8v1-A (Illumina) | 951117 | 3 | 0 | - | - |
| Total |  |  | 9002 | 1551 | 8802 | 1519 |

(a) Number of participants in dataset with neuroimaging scans available. (b) Participants with genotype call rate less than 95%. Genotype call rates for participants in other datasets with neuroimaging scans was at least 95%. PC scores were not evaluated for datasets without any participants with neuroimaging scans available.

**Figure S1:**
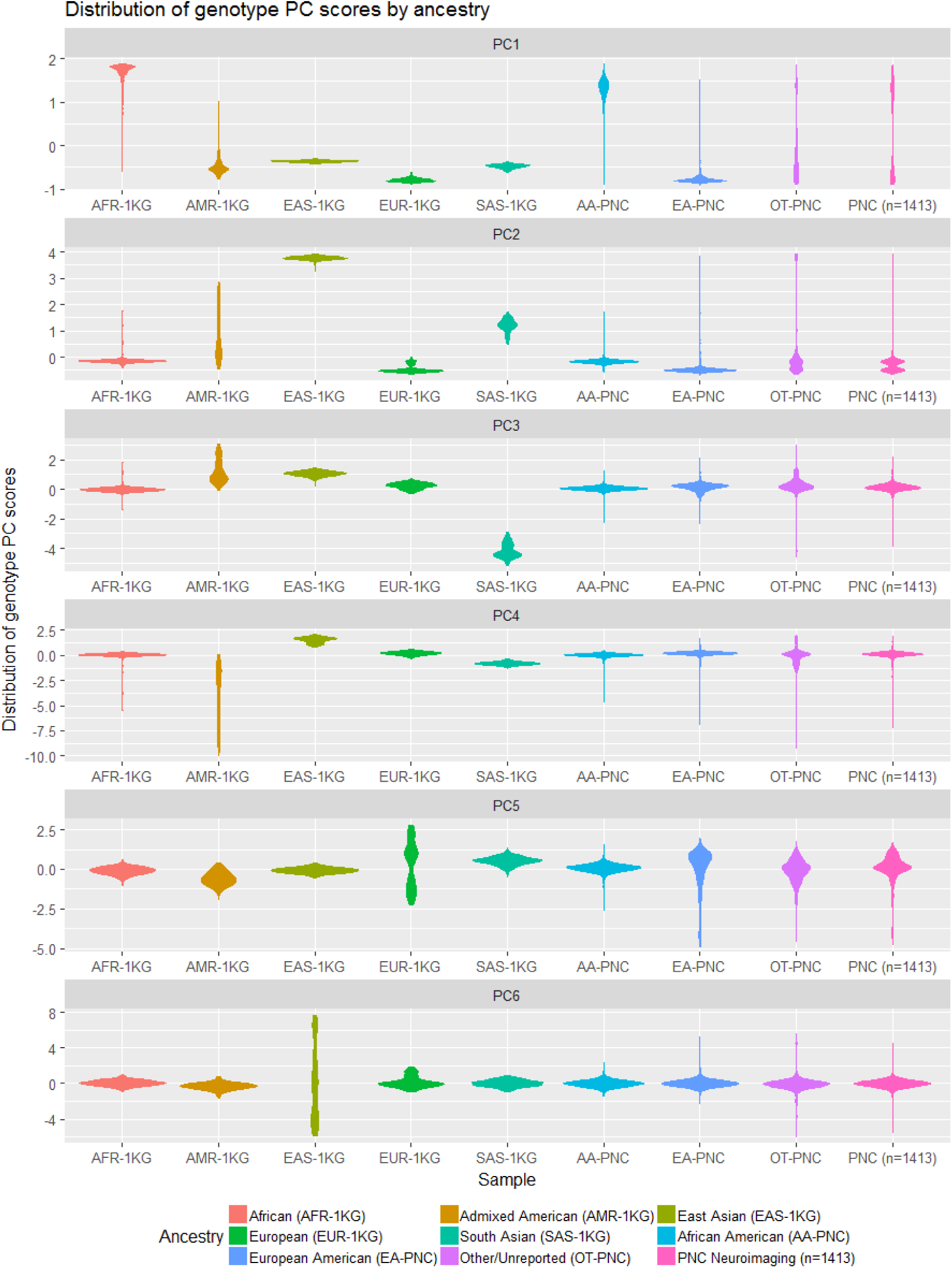
Distributions of scores for first six genotype principal components by ancestry groups and PNC neuroimaging sample (n=1,413). PCs are from genetic relatedness matrix of genotype sample from PNC (n=8,802 after quality control filters) and reference sample from 1000 Genomes (1KG) phase 3 project (n=2,504). Violin plots indicate distributions of PC scores by five ancestry groups in 1KG project (AFR-1KG, AMR-1KG, EAS-1KG, EUR-1KG, and SAS-1KG); self-reported African Americans (AA-PNC), European Americans (EA-PNC), and participants reporting neither race (OT-PNC) in PNC’s genotype sample (n=8,802); and the neuroimaging subsample of the PNC (n=1,413) studied in the analysis.

**Figure S2:**
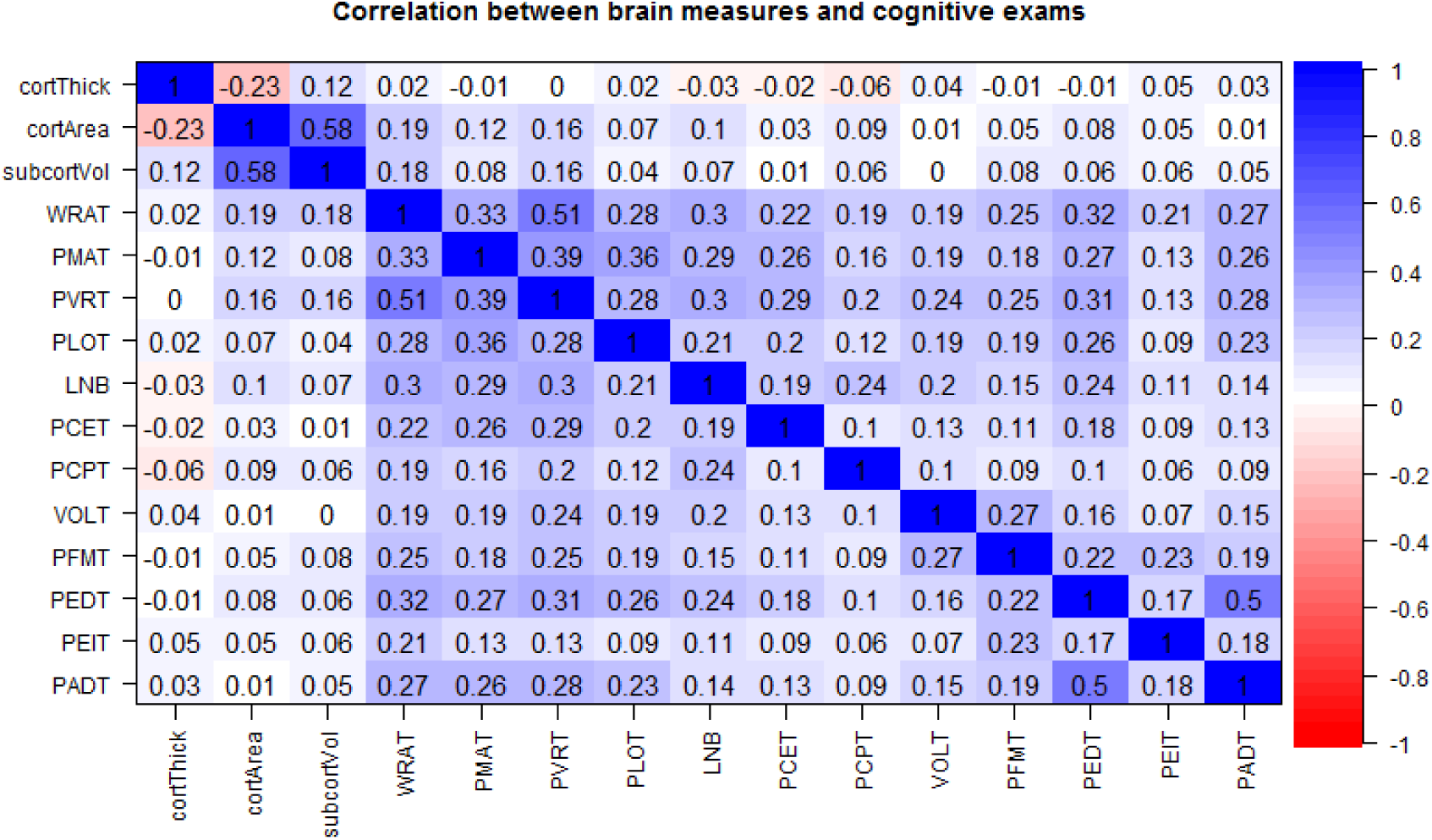
Correlations between brain measures and cognitive exams in the PNC (n=1,413). Prior to computing correlations, each measure was residualized against age, sex, and first six genotype PCs.

**Figure S3:**
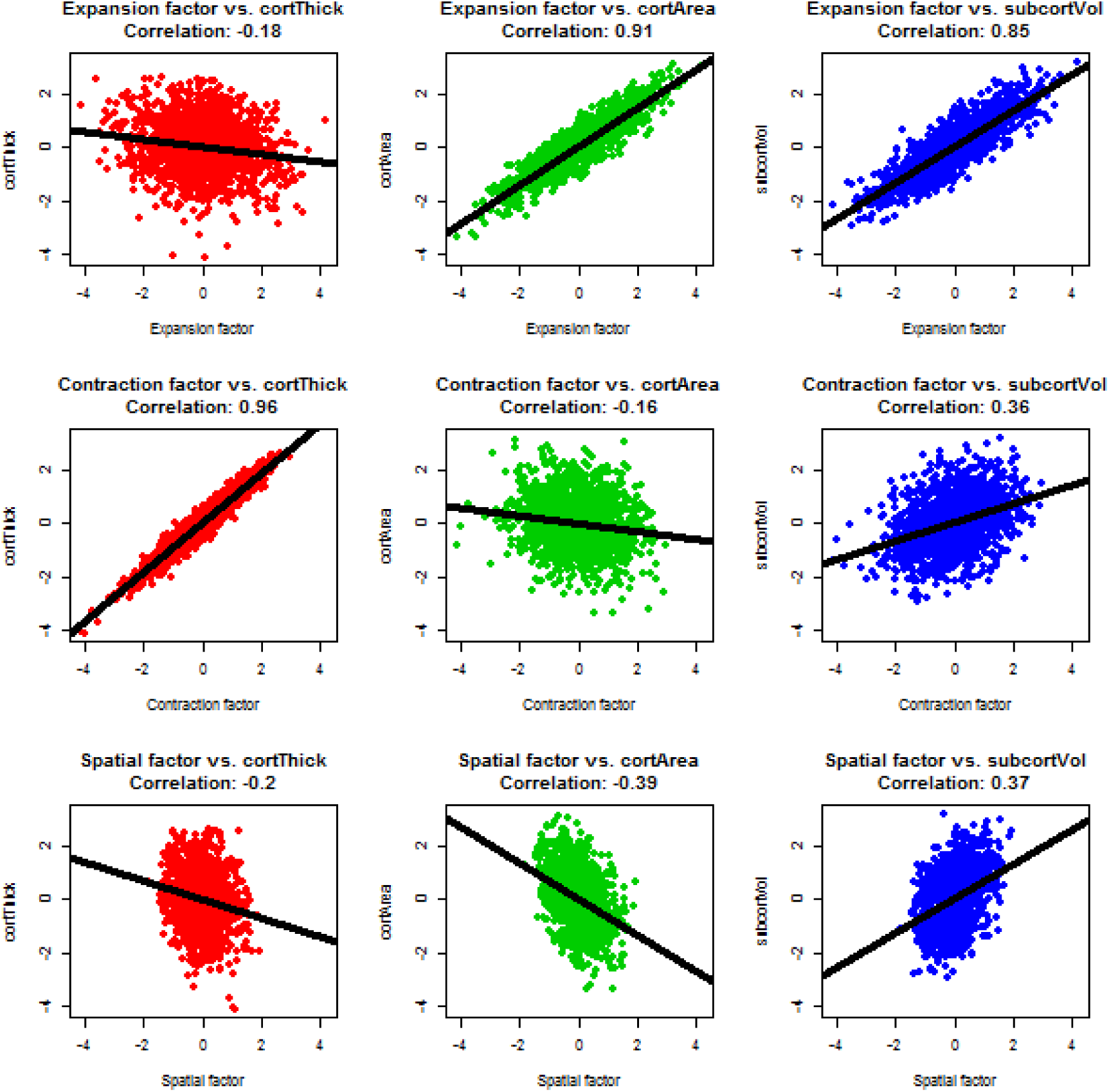
Pairwise scatter plots between three neurodevelopmental factors and brain measures residualized against age, sex, and first six genotype PCs and then scaled to zero mean and unit variance.

